# Elexacaftor/Tezacaftor/Ivacaftor (Trikafta^®^) does not affect the Growth of *Aspergillus fumigatus in vitro*

**DOI:** 10.1101/2024.08.12.607675

**Authors:** Elise Biquand, Sandra Khau, Delphine Fouquenet, Floriane Costes, Adélaïde Chesnay, Guillaume Desoubeaux, Benoit Briard

## Abstract

Fungal infection, especially allergic bronchopulmonary aspergillosis, is a leading cause of infection-associated morbidity and death in patients with cystic fibrosis. We have previously discovered that the new leading therapeutic treatment in cystic fibrosis, CFTR modulators Elexakaftor/Tezacaftor/Ivacaftor (ETI, Trikafta^®^), drastically reduces the colonization and infection by *Aspergillus fumigatus*. However, the reasons for this decrease in patients are not known so far. In this study we have shown, using *A. fumigatus* reference strains and strains isolated from patients with cystic fibrosis, that CFTR modulators have no discernable impact on fungal growth *in vitro*. These results are consistent regardless of drugs concentration, demonstrating that treatment does not exert a direct antimicrobial or stimulatory effect *in vitro* on *A. fumigatus* conidia growth, when exposed to actual *in vivo* CFTR modulators concentration.

## Importance

The advent of ETI therapy represents a pivotal moment, signaling the onset of major changes in the medical field of cystic fibrosis and its related infectious diseases. However, impact of ETI treatment on patient’s microbiota and pathogens has to be further studied as proofs arise of changes in patient colonization.

## Observation

We previously published pioneering data that underscored a substantial change in the impact of *Aspergillus*-related diseases in patients with cystic fibrosis (pwCF) treated with a combination of CFTR (cystic fibrosis transmembrane conductance regulator) modulators (1). Our results highlighted a noticeable reduction in the proportion of respiratory samples colonized by *Aspergillus* spp. Additionally, there was a statistically-significant decrease in anti-*Aspergillus* precipitin reactivity and in total IgE concentration following the initiation of Trikafta^®^ treatment. These findings suggested a robust decline in fungal development and colonization. Several mechanisms could be raised to explain such outcomes. For instance, ETI could suppress the overall inflammatory response. Indeed, by reducing the neutrophil activation, by moderating the release of reactive oxygen species (ROS) and cytokines, and by restoring the macrophage-induced phagocytosis (2), the therapeutic combination can make the local environment more challenging for microbial colonization and development of immune hyperreactivity. Furthermore, next-generation sequencing (NGS) revealed that CFTR modulators enhance microbial diversity in respiratory samples, restoring the steady state of the microbiota: major colonizer pathogens being less represented, while minor species regained their prominence (3). For instance, lung colonization by *Pseudomonas aeruginosa* bacillus was shown to be decreased by 24.2% in 124 patients receiving ETI for one year (4).

However, the precise mechanism of action of CFTR modulators remains elusive in such a context. It is acknowledged that some non-anti-infective drugs can *in vitro* exert a direct antimicrobial effect against pathogens: *e*.*g*. mycophenolate mofetil – conventionally used for the prevention of graft rejection - presents low minimal inhibitory concentrations (MICs) against various fungal species (5) and might play a critical role in *in vivo* strain selection (6).

To investigate this, we assessed fungal growth from the conidial stage of two reference strains (DAL and Af293) and four clinical isolates (two issued from ETI-treated patients, and two from non-treated ones), in different conditions with or without ETI treatment. Several media were exploited for the *in vitro* cultures: Dulbecco’s modified Eagle medium (DMEM), Roswell Park Memorial Institute medium (RPMI), as well as minimal medium (MM) which is a defined medium classically used to test compounds acting on *A. fumigatus*. ETI drugs were tested at two different concentrations: serum concentrations ([3VX]_S_) based on clinical observations from therapeutic drug monitoring in treated patients (9.5 μM VX-445, 6.53 μM VX-661 and 2.29 μM VX-770) (7); and effective concentrations ([3VX]_E_) established according to *in vitro* efficacy data on epithelial cells to restore CFTR channel activity (3 μM VX-445, 18 μM VX-661 and 1 μM VX-770) (8).

Regardless of the combination of concentrations ([3VX]_S_ and [3VX]_E_) or the medium used, the growth of *A. fumigatus* DAL strain was not perturbed by ETI treatment (Figure 1A, B). Similarly, fungal biofilm development remained unaffected by the treatment (Figure 1C, and video in Supplementary Data 1). Interestingly none of the clinical strains obtained from pwCF who were either treated or untreated with Elexacaftor/Tezacaftor/Ivacaftor exhibited susceptibility to the treatment compared to the vehicle condition (Figure 2).

**Figure 1.**
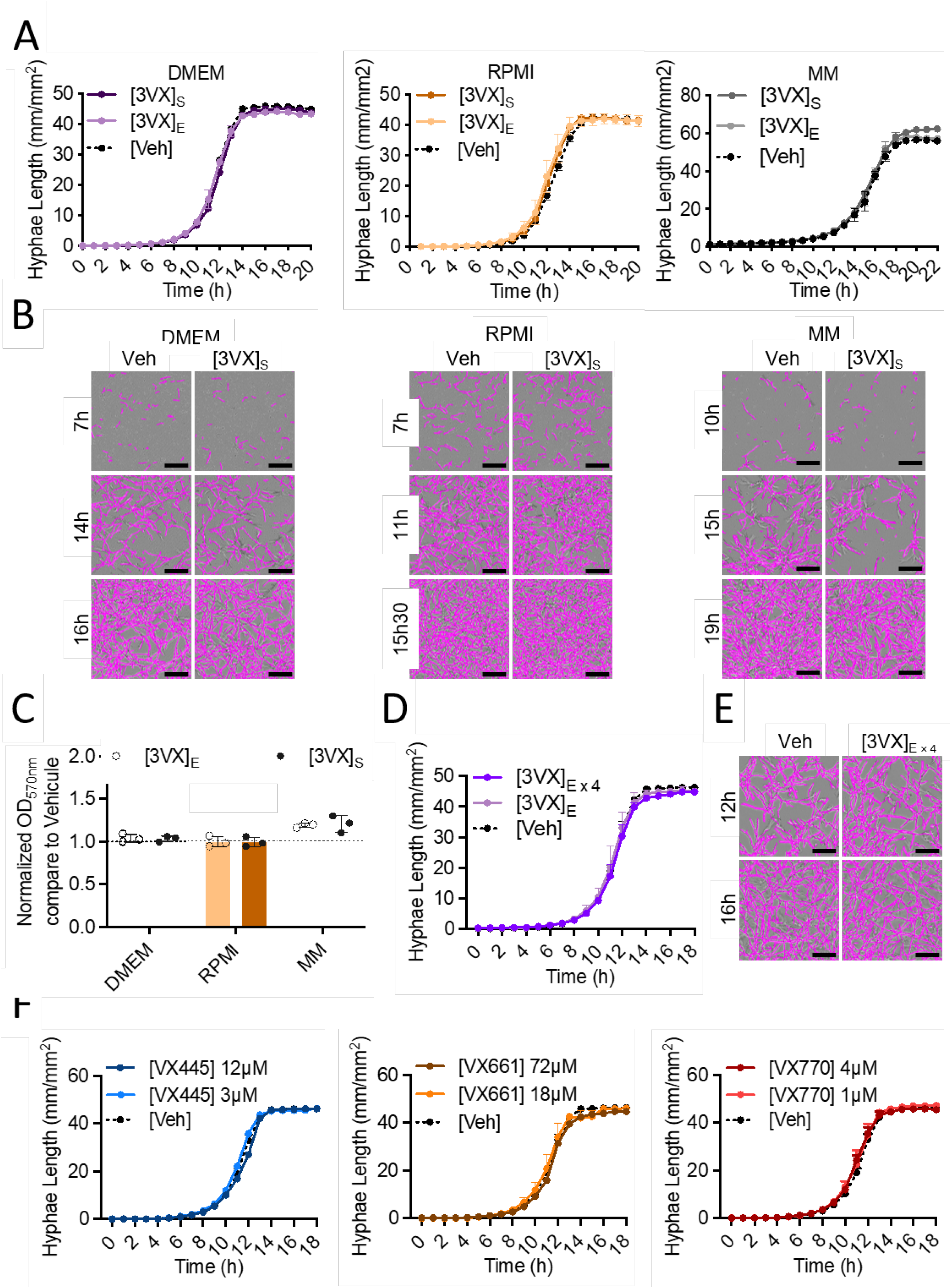
Evaluation of the direct antimicrobial effects of CFTR modulators (elexacaftor, VX445 / tezacaftor, VX661 / ivacaftor, VX770) on *Aspergillus fumigatus* growth and biofilm formation. **A** – Measurement of *A. fumigatus* hyphal length (Strain: DAL) using the IncuCyte^®^ live-cell analysis system in DMEM and RPMI media supplemented with 10% fetal bovine serum (FBS) or minimal medium (MM) in the presence of 3VX (VX445 / VX661 / VX770) treatment at effective concentration ([3VX]_E_) or the serum concentration found in treated cystic fibrosis patients ([3VX]_S_), as well as with vehicle ([Veh]; DMSO). The shown data depict a representative experiment; the experiment was independently replicated at least 3 times, with measurements taken hourly in each well. **B** – Representative images of *A. fumigatus* (Strain: DAL) biofilm at indicated times and media with or without 3VX treatment at patient serum concentration. Fungal hyphae were detected using the neurotrack module (Sartorius) and highlighted in purple. Scale bars, 200 μm. **C** – Assessment of biofilm formation (Strain: DAL) by crystal violet testing and endpoint measurement at 570nm after culture in DMEM and RPMI media supplemented with 10% FBS or MM (*n*=3 for each condition). **D** – Measurement of hyphal length of *A. fumigatus* (Strain: DAL) in DMEM with 10% FBS, with or without 3VX treatment at effective concentration or 4-fold effective concentration. **E** – Representative images of *A. fumigatus* (Strain: DAL) biofilm at 12 and 16 h in DMEM with or without 3VX treatment at 4-fold effective concentration. Scale bars, 200 μm. **F** – Measurement of hyphal length in DMEM with 10% FBS with VX445, VX661, or VX770 at the effective concentration or 4-fold effective concentration. Data are shown as mean ± SD. Representative experiment shown for A, B, D – F.

**Figure 2.**
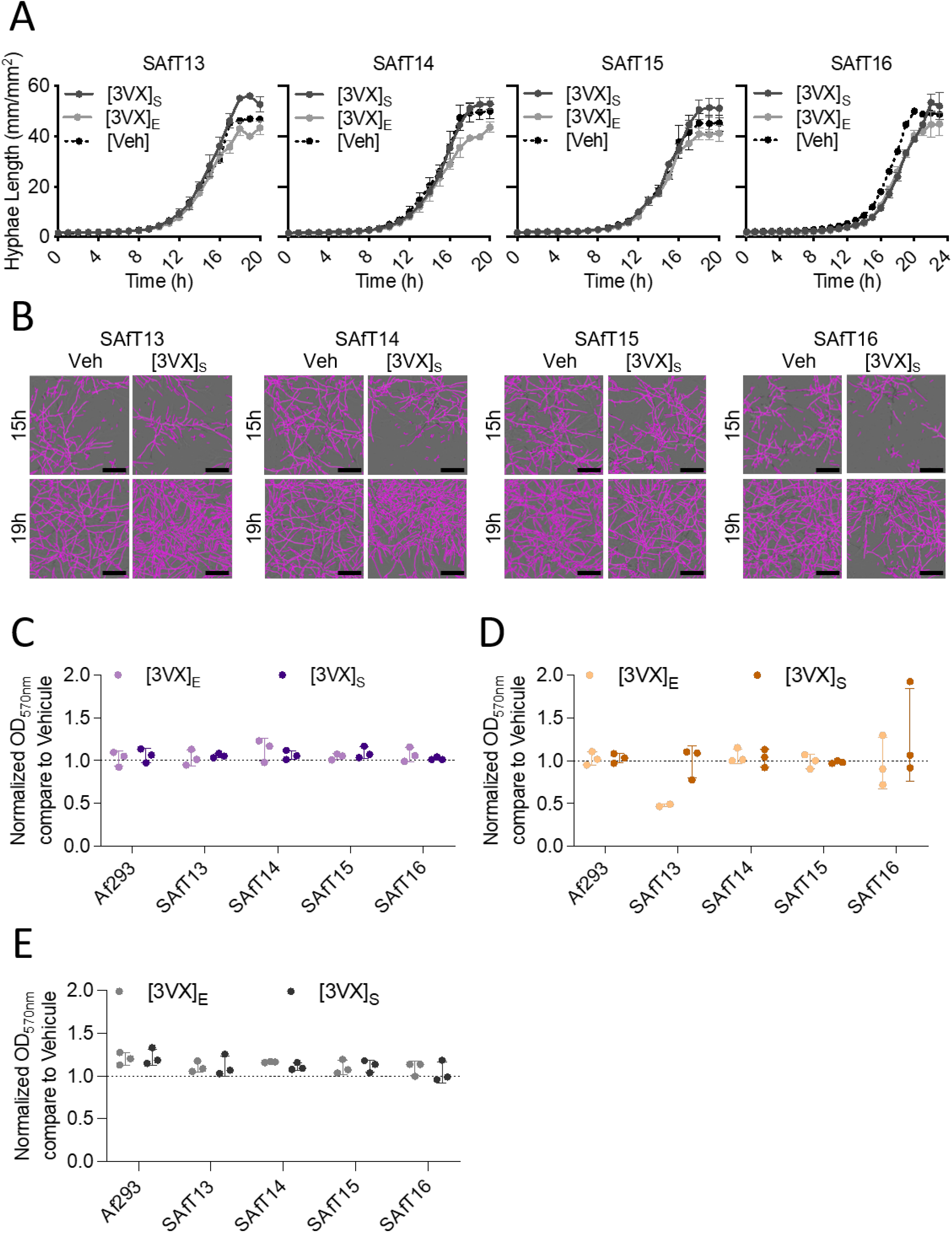
Evaluation of the direct antimicrobial effects of CFTR modulators on clinical *Aspergillus fumigatus* strains derived from cystic fibrosis patients. Four clinical strains from cystic fibrosis patients (SAfT13 to SAfT16) were used. **A** – Measurement of *A. fumigatus* hyphal length using the IncuCyte^®^ live-cell analysis system in minimal medium (MM) in the presence of 3VX (VX445 / VX661 / VX770) treatment at effective concentration ([3VX]_E_) or the serum concentration found in treated cystic fibrosis patients ([3VX]_S_), as well as with vehicle ([Veh]; DMSO). The shown data depict a representative experiment; the experiment was independently replicated at least 3 times, with measurements taken hourly in each well. **B** – Representative images of *A. fumigatus* biofilm at indicated times with or without 3VX treatment at patient serum concentration. Fungal hyphae were detected using the neurotrack module (Sartorius) and highlighted in purple. Scale bars, 200 μm. **C**-**E** – Assessment of biofilm formation by crystal violet testing and endpoint measurement at 570nm after culture in DMEM (**C**) and RPMI (**D**) media supplemented with 10% FBS or MM (**E**) in the presence of 3VX treatment at effective concentration ([3VX]_E_) or the serum concentration observed in treated cystic fibrosis patients ([3VX]_S_), as well as with vehicle ([Veh]; DMSO). Testing included one reference strain (Af293) and four clinical strains from cystic fibrosis patients (SAfT13 to SAfT16). Normalization was carried out based on the vehicle control. Data are presented as mean ± SD, *n*=3.

Furthermore, despite a significant increase (4-fold) in the combined concentration of all three ETI, no effects were observed on the growth of *A. fumigatus* (Figure 1D, E, and video in Supplementary Data 2). We wondered whether the three compounds would exhibit activity when used separately on *A. fumigatus*. We tested each compound at two different concentrations, corresponding to the effective concentration or a 4-fold increase. However, at all concentrations tested, the growth *of A. fumigatus* was not affected by any of the compounds compared to vehicle condition (Figure 1F).

Considering our results, it is plausible to assert that the ETI combination does not exert a direct antimicrobial or stimulatory effect *in vitro* on *A. fumigatus* conidia, when exposed to actual *in vivo* ETI concentrations. Interestingly, Jones JT *et al*. (9) recently showed on a preformed biofilm that at 5μM CFTR modulators were able to dramatically reduce fungal biomass by 50%, increase cell permeability close to amphotericin B activity, and raise metabolic activity of *A. fumigatus* by 20-30% (9). Therefore, we decided to increase the compound concentration to 5 μM for each and test the effect on resting conidia growth in RPMI, DMEM, or MM media. Unfortunately, no significant effect was observed on fungal growth compared to the vehicle control (Supplementary Data 3). However, this previous work was performed on the preform hyphal stage, which may explain the difference observed (9). As patients present lower serum concentration of modulators and are infected by conidia, the effect of Trikafta^®^ treatment on *A. fumigatus* growth is presumably indirect. As it has been shown that Trikafta^®^ treatment sensitize *Aspergillus* to antifungal drugs (9), it would be interesting to assess if the fungus is more sensitive to immune cells or antimicrobial peptides secreted by the host. Such increased sensitivity could participate on the observed reduction of *Aspergillus* infections in patients (1). Thus, the introduction of ETI therapy marks a crucial milestone, heralding the start of significant transformations in the medical landscape for cystic fibrosis and related infectious diseases but more mechanistic investigations on modulation of microbiota and mycobiota are still needed.

## Material & Methods

### Aspergillus strains and culture

Two *Aspergillus fumigatus* reference strains were used, DAL (CEA10, CBS144.89) and Af293 (NCPF7367), and four clinical isolates (SAfT14 & SAfT16 issued from ETI-treated patients, and SAfT13 & SAfT15 from non-treated patients). *A. fumigatus* strains were grown on 2% (w/v) malt/2% (w/v) agar slants for 1 week. Conidia were harvested in water containing 0.1% (v/v) Tween-20 and filtered through cell strainer 40μM (Fisherbrand). *In vitro* cultures were realized in three media: Dulbecco’s modified Eagle medium (DMEM, Gibco) and Roswell Park Memorial Institute medium (RPMI, Gibco), both supplemented with 10% fetal bovine serum (FBS), as well as minimal medium (MM)(10). ETI drugs (Ivacaftor (VX-770), Tezafactor (VX-661) and Elexacaftor (VX-445) (MedChemExpress, HY-13017, HY-15448 & HY-111772)) were mixed and tested at two different concentrations: serum concentrations ([3VX]_S_ = 9.5 μM VX-445, 6.53 μM VX-661 and 2.29 μM VX-770); and effective concentrations ([3VX]_E_ = 3 μM VX-445, 18 μM VX-661 and 1 μM VX-770). Vehicle suspension (DMSO) was used as negative control.

### Growth assay

The level of fungal growth of 1.10^3^ *A. fumigatus* conidia, deposited in wells of a 96w-plate, was continuously monitored at 37°C by an IncuCyte^®^ live-cell analysis system (Sartorius™, Göttingen – Germany) for 20h in DMEM+10% FBS and RPMI+10% FBS, or 24h for MM. Hyphae length was analysed using the NeuroTrack module of IncuCyte (11). At endpoint, the biofilm was stained with 0.01% crystal-violet, washed two times and discolored with 30% acetic acid, to measure absorbance at 570nm with the Multiskan Sky spectrophotometer (Thermo Fisher Scientific™, Waltham, MA – USA) (12).

## Author contribution

BB and EB conceived the concept. EB, SK, DF and FC carried out the experiments. AC supervised the study. GD, BB and EB led the writing. All the authors edited the text and approved the final version of the manuscript.

## Acknowledgment

The authors thank Dr. Mustapha Si-Tahar for his valuable support. The authors are grateful to the department of medical mycology of Tours university hospital (France) for collecting the clinical strains that were used in the experiments.

## Figure legend

**Supplementary Data 1**. Video illustrating the biofilm formation of *A. fumigatus* in DMEM (**A**), RPMI (**B**), and MM (**C**), assessed by Incucyte (Sartorius), with and without 3VX treatment at serum concentrations observed in treated cystic fibrosis patients ([3VX]_S_). The fungal hyphae are detected using the neurotrack module (Sartorius) and highlighted in purple. Scale bars, 100 μm.

**Supplementary Data 2:** Video illustrating the biofilm formation of *A. fumigatus* (Strain: DAL), assessed by Incucyte (Sartorius), with and without 3VX treatment at 4-fold serum concentrations ([3VX]_E x 4_) in DMEM supplemented with 10% FBS. The fungal hyphae are detected using the neurotrack module (Sartorius) and highlighted in purple. Scale bars, 100 μm.

**Supplementary Data 3**. Evaluation of the direct antimicrobial effects of CFTR modulators at 5μM on *Aspergillus fumigatus DAL* strain. Measurement of *A. fumigatus* hyphal length using the IncuCyte^®^ live-cell analysis system in (**A**) DMEM, (**B**) RPMI or (**C**) minimal medium (MM) in the presence of 5μM of 3VX (VX445 / VX661 / VX770) treatment, as well as with vehicle ([Veh]; DMSO). The shown data depict a representative experiment; the experiment was independently replicated 3 times, with measurements taken hourly in each well. **D** – Assessment of biofilm formation by crystal violet testing and endpoint measurement at 570nm after culture in DMEM and RPMI media supplemented with 10% FBS or MM in the presence of 3VX treatment at 5μM each compound as well as with vehicle ([Veh]; DMSO). Data are presented as mean ± SD, *n*=3.

## References

1. Chesnay A, Bailly É, Cosson L, Flament T, Desoubeaux G. 2022. Advent of elexacaftor/tezacaftor/ivacaftor for cystic fibrosis treatment: What consequences on Aspergillus-related diseases? Preliminary insights. J Cyst Fibros 21:1084–1085.

2. Bercusson A, Jarvis G, Shah A. 2021. CF fungal disease in the age of CFTR modulators. Mycopathologia 186:655–664.

3. Hong G, Daniel SG, Lee J-J, Bittinger K, Glaser L, Mattei LM, Dorgan DJ, Hadjiliadis D, Kawut SM, Collman RG. 2023. Distinct community structures of the fungal microbiome and respiratory health in adults with cystic fibrosis. J Cyst Fibros 22:636–643.

4. Beck MR, Hornick DB, Pena TA, Singh SB, Wright BA. 2023. Impact of elexacaftor/tezacaftor/ivacaftor on bacterial cultures from people with cystic fibrosis. Pediatr Pulmonol 58:1569–1573.

5. Oz HS, Hughes WT. 1997. Novel anti-Pneumocystis carinii effects of the immunosuppressant mycophenolate mofetil in contrast to provocative effects of tacrolimus, sirolimus, and dexamethasone. J Infect Dis 175:901–904.

6. Hoffmann CV, Nevez G, Moal M-C, Quinio D, Le Nan N, Papon N, Bouchara J-P, Le Meur Y, Le Gal S. 2021. Selection of Pneumocystis jirovecii inosine 5’-monophosphate dehydrogenase mutants in solid organ transplant recipients: implication of mycophenolic acid. J Fungi Basel Switz 7:849.

7. Pigliasco F, Cafaro A, Stella M, Baiardi G, Barco S, Pedemonte N, D’Orsi C, Cresta F, Casciaro R, Castellani C, Calevo MG, Mattioli F, Cangemi G. 2023. Simultaneous Quantification of Ivacaftor, Tezacaftor, and Elexacaftor in Cystic Fibrosis Patients’ Plasma by a Novel LC-MS/MS Method. Biomedicines 11:628.

8. Becq F, Mirval S, Carrez T, Lévêque M, Billet A, Coraux C, Sage E, Cantereau A. 2022. The rescue of F508del-CFTR by elexacaftor/tezacaftor/ivacaftor (Trikafta) in human airway epithelial cells is underestimated due to the presence of ivacaftor. Eur Respir J 59:2100671.

9. Jones JT, Morelli KA, Vesely EM, Puerner CTS, Pavuluri CK, Ross BS, van Rhijn N, Bromley MJ, Cramer RA. 2023. The cystic fibrosis treatment Trikafta affects the growth, viability, and cell wall of Aspergillus fumigatus biofilms. mBio 14:e0151623.

10. Cove DJ. 1966. The induction and repression of nitrate reductase in the fungus Aspergillus nidulans. Biochim Biophys Acta 113:51–56.

11. Wurster S, Sass G, Albert ND, Nazik H, Déziel E, Stevens DA, Kontoyiannis DP. 2020. Live imaging and quantitative analysis of Aspergillus fumigatus growth and morphology during inter-microbial interaction with Pseudomonas aeruginosa. Virulence 11:1329–1336.

12. Briard B, Bomme P, Lechner BE, Mislin GLA, Lair V, Prévost M-C, Latgé J-P, Haas H, Beauvais A. 2015. Pseudomonas aeruginosa manipulates redox and iron homeostasis of its microbiota partner Aspergillus fumigatus via phenazines. Sci Rep 5:8220.

